# Application of transposon-insertion sequencing to determine gene essentiality in the acetogen *Clostridium autoethanogenum*

**DOI:** 10.1101/2021.05.19.444907

**Authors:** Craig Woods, Christopher M. Humphreys, Claudio Tomi-Andrino, Anne M. Henstra, Michael Köpke, Sean D. Simpson, Klaus Winzer, Nigel P. Minton

**Affiliations:** Clostridia Research Group, BBSRC/EPSRC Synthetic Biology Research Centre (SBRC), Biodiscovery Institute, School of Life Sciences, The University of Nottingham, Nottingham, NG7 2RD, UK; Centre for Analytical Bioscience, Advanced Materials and Healthcare Technologies Division, School of Pharmacy, The University of Nottingham, Nottingham, NG7 2RD, UK; BBSRC/EPSRC Synthetic Biology Research Centre (SBRC), School of Mathematical Sciences, The University of Nottingham, Nottingham, NG7 2RD, UK; LanzaTech Inc., 8045 Lamon Avenue, Suite 400, Skokie, IL, USA; NIHR Nottingham Biomedical Research Centre, Nottingham University Hospitals NHS Trust and the University of Nottingham, Nottingham, NG7 2RD, UK

## Abstract

The majority of the genes present in bacterial genomes remain poorly characterised with up to one third of those that are protein encoding having no definitive function. Transposon insertion sequencing represents a high-throughput technique that can help rectify this deficiency. The technology, however, can only be realistically applied to easily transformable species leaving those with low DNA-transfer rates out of reach. Here we have developed a number of approaches that overcome this barrier in the autotrophic species *Clostridium autoethanogenum* using a *mariner-based* transposon system. The inherent instability of such systems in the *Escherichia coli* conjugation donor due to transposition events was counteracted through the incorporation of a conditionally lethal *codA* marker on the plasmid backbone. Relatively low frequencies of transformation of the plasmid into *C. autoethanogenum* were circumvented through the use of a plasmid that is conditional for replication coupled with the routine implementation of an Illumina library preparation protocol that eliminates plasmid-based reads. A transposon library was then used to determine the essential genes needed for growth using carbon monoxide as a sole carbon and energy source.

**IMPORTANCE:** Although microbial genome sequences are relatively easily determined, assigning gene function remains a bottleneck. Consequently, relatively few genes are well characterised, leaving the function of many as either hypothetical or entirely unknown. High-throughput, transposon sequencing can help remedy this deficiency, but is generally only applicable to microbes with efficient DNA-transfer procedures. These exclude many microorganisms of importance to humankind either as agents of disease or as industrial process organisms. Here we developed approaches to facilitate transposon-insertion sequencing in the acetogen *Clostridium autoethanogenum*, a chassis being exploited to convert single-carbon waste gases, CO and CO_2_, into chemicals and fuels at an industrial scale. This allowed the determination of gene essentiality under heterotrophic and autotrophic growth providing insights into the utilisation of CO as a sole carbon and energy source. The strategies implemented are translatable and will allow others to apply transposon-insertion sequencing to other microbes where DNA-transfer has until now represented a barrier to progress.

## INTRODUCTION

Although microbial genome sequences are relatively easily determined, assigning gene function remains a bottleneck. Consequently, relatively few genes are well characterised, leaving the function of many as either hypothetical or entirely unknown. Thus, even the Syn 3.0 minimal genome retains 149 genes (32%) of unknown function ^1^. A greater understanding of gene functionality can be gleaned through the deployment of high throughput transposon sequencing. This technique is characterised by the simultaneous Illumina sequencing of the site of transposon insertion in pooled mutant libraries using a sequencing primer specific to the transposon-chromosomal junction. If the library consists of a sufficiently high number of unique insertions then the required gene set for the growth conditions used can be inferred since unrepresented or underrepresented genes are likely to be essential. There are several names for this type of approach, including the first four all published in 2009: TraDIS ^2^, HITS ^3^, Tn-Seq and INSeq ^4^. All of these techniques aim to identify the position and quantity of transposon mutants and are collectively referred to as Transposon insertion sequencing (TIS) ^5^.

The deployment of TIS typically is largely dependent on high frequency DNA transfer. This excludes its application to many microbial species. Anaerobic bacteria, and in particular members of the genus *Clostridium*, are of both medical and industrial importance but generally display low rates of DNA transfer. This has limited the exploitation of TIS in this grouping where to date TIS has only been applied ^6^ to the pathogen *Clostridioides difficile* (formerly *Clostridium difficile*). One group of bacteria with increasing importance are the anaerobic acetogens, typified by *Clostridium autoethanogenum*. Acetogens possess the Wood-Ljungdahl pathway (WLP), or reductive acetyl-CoA pathway, which allows the fixation of CO and CO_2_ ^7^. Suggested to be the earliest autotrophic pathway ^8^, it is the most energy efficient of the seven known carbon fixation pathway since it conserves energy while all others require its input ^9^. Reducing equivalents needed for metabolic processes are obtained either from H_2_ or CO using hydrogenases or CO dehydrogenase (CODH), respectively. Carbon is fixed via the Eastern branch of the pathway where, through a series of cobalamin and tetrahydrofolate-dependent reactions, CO_2_ is reduced to a methyl group. The methyl group from the Eastern branch is then combined with CO to form acetyl-CoA which is the root of subsequent anabolic reactions ^10-13^.

While the majority of acetogens synthesize acetate as the sole fermentation product some, typified by *C. autoethanogenum*, naturally produce industrially relevant compounds as 2-3 butanediol and ethanol, the latter on a commercial scale ^14^. Commercial efforts to extend the product range further are ongoing with isopropanol being a notable example ^15^. *C. autoethanogenum* is one of the best understood autotrophic acetogens with a manually annotated genome ^16,17^ and has been subjected to transcriptomic and proteomic analysis ^18^.

In the current study we sought to maximise the benefit of available *C. autoethanogenum* genome data through implementation of TIS. However, as DNA transfer into *C. autoethanogenum* is only possible at relatively low frequencies, a number of essential modifications to the procedure were required. Specifically, the use of a conditional replicon and an inducible orthogonal expression system to control production of transposase allowed the controlled generation of a large mutant library from a small number of initial transconjugant colonies. Additionally, the incorporation of I-SceI recognition sequences into the delivery vehicle provided a mechanism to eliminate those mini-transposon sequences still present on autonomous copies of the plasmid during the transposon mutant library preparations stage. These adaptations have allowed a thorough genetic analysis of the WLP in *C. autoethanogenum* and, for the first time, the determination of the essential gene set required for growth on CO as a sole carbon and energy source.

## RESULTS

### Control of transposition

A fundamental requirement of an effective transposon-delivery system is that transposition should preferentially take place in the target strain and not in the donor strain. This may be achieved through the use of an orthogonal expression system in which the promoter controlling production of the transposase is recognised by transcriptional factors that are only present in the target microbe. A previously described clostridial system ^19^ exploited the *Clostridioides difficile* alternate sigma factor TcdR ^19^ and one of the only two promoters it recognises, the P_*tcdB*_ promoter of the toxin B gene (*tcdB*). By generating a derivative of *Clostridium acetobutylicum* in which the TcdR-encoding *tcdR* gene was inserted into the genome at the *pyrE* locus, any subsequently introduced gene that was placed under the control of the P_*tcdB*_ promoter is expressed. In this example, the gene placed under the control of this orthogonal system was the mariner transposase gene ^19^.

For exploitation in *C. autoethanogenum*, further control was engineered into the system by placing expression of *tcdR* (Fig. 1) under the control of a lactose inducible promoter *P*_*bgaL*_ previously shown to be functional in the closely related *Clostridium ljungdahlii* ^20^. Accordingly, the *P*_*bgaL*_ promoter, together with the necessary *bgaR* which encodes a transcriptional regulator, was positioned 5′ to the *tcdR* gene and the DNA module created (*bgaR*-P_*bgaL*_::*tcdR*) integrated into the *C. autoethanogenum* chromosome at the *pyrE* locus using ACE (Allele Coupled Exchange) ^20-23^. This involved restoring a uracil-requiring Δ*pyrE* mutant strain to prototrophy concomitant with genomic insertion of *bgaR*-P_*gaL*_::*tcdR* using the ACE plasmid pMTL-CH20lactcdR. Successful mutant generation was confirmed by PCR analysis and Sanger sequencing of the amplified DNA and the resulting strain was termed *C. autoethanogenum* C24.

**Figure 1.**
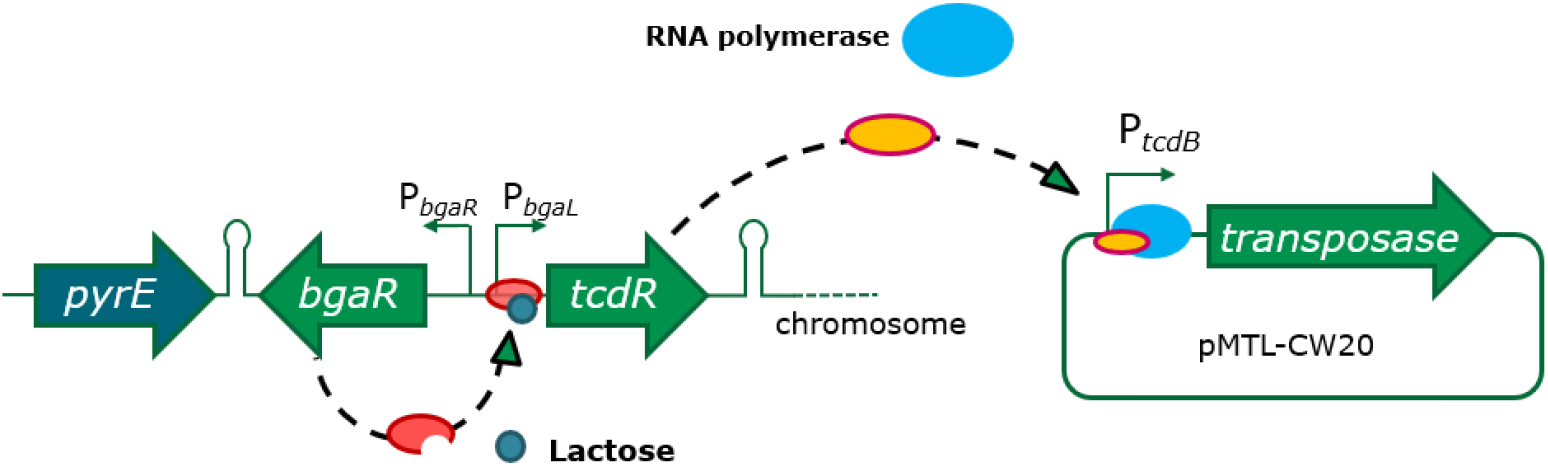
TcdR-mediated orthogonal expression. In *C. autoethanogenum* C24, *tcdR* is under the control of the lactose-inducible promoter system *bgaR*-P_*bgaL*_ from *C. perfringens*. In this way the P_*tcdB*_ promoter can be induced indirectly via the inducible expression of *tcdR* from the chromosome.

To confirm that TcdR production could be controlled by the addition of exogenous lactose in strain C24, the *Clostridium perfringens catP* reporter gene encoding a chloramphenicol acetyltransferase (CAT) was cloned downstream of the P_*tcdB*_ promoter on an appropriate clostridial shuttle vector. Regulation of the reporter gene was shown to be dependent on the addition of the lactose inducer (Figure 2). We therefore chose to use this expression system to create our transposon library by placing the transposase under the control of P_*tcdB*_ on the transposon-delivery plasmid pMTL-CW20 (Fig. 3).

**Figure 2.**
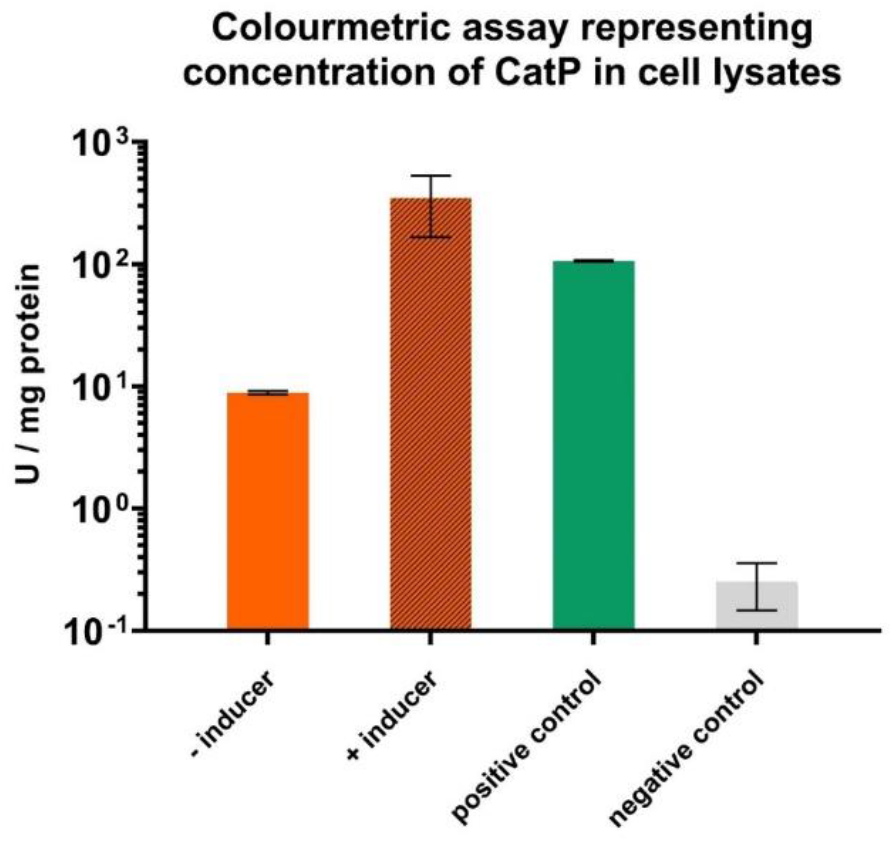
Chloramphenicol acetyl transferase (CAT) assay of lactose-inducible orthogonal system. Expression from P_*tcdB*_ was quantified using a CAT assay. Three plasmids were conjugated into *C. autoethanogenum P*_*bgaL*_*_tcdR* (*C. autoethanogenum* C24) with each plasmid harbouring *catP* under the control of either P_*tcdB*_, P_*thl*_ (positive control), or no promoter (negative control). The strain harbouring the P_*tcdB*_ plasmid was tested with and without the addition of 10 mM lactose while the remaining plasmids were tested without lactose. The data shown is the result of biological triplicates with error bars showing the standard deviation.

**Figure 3.**
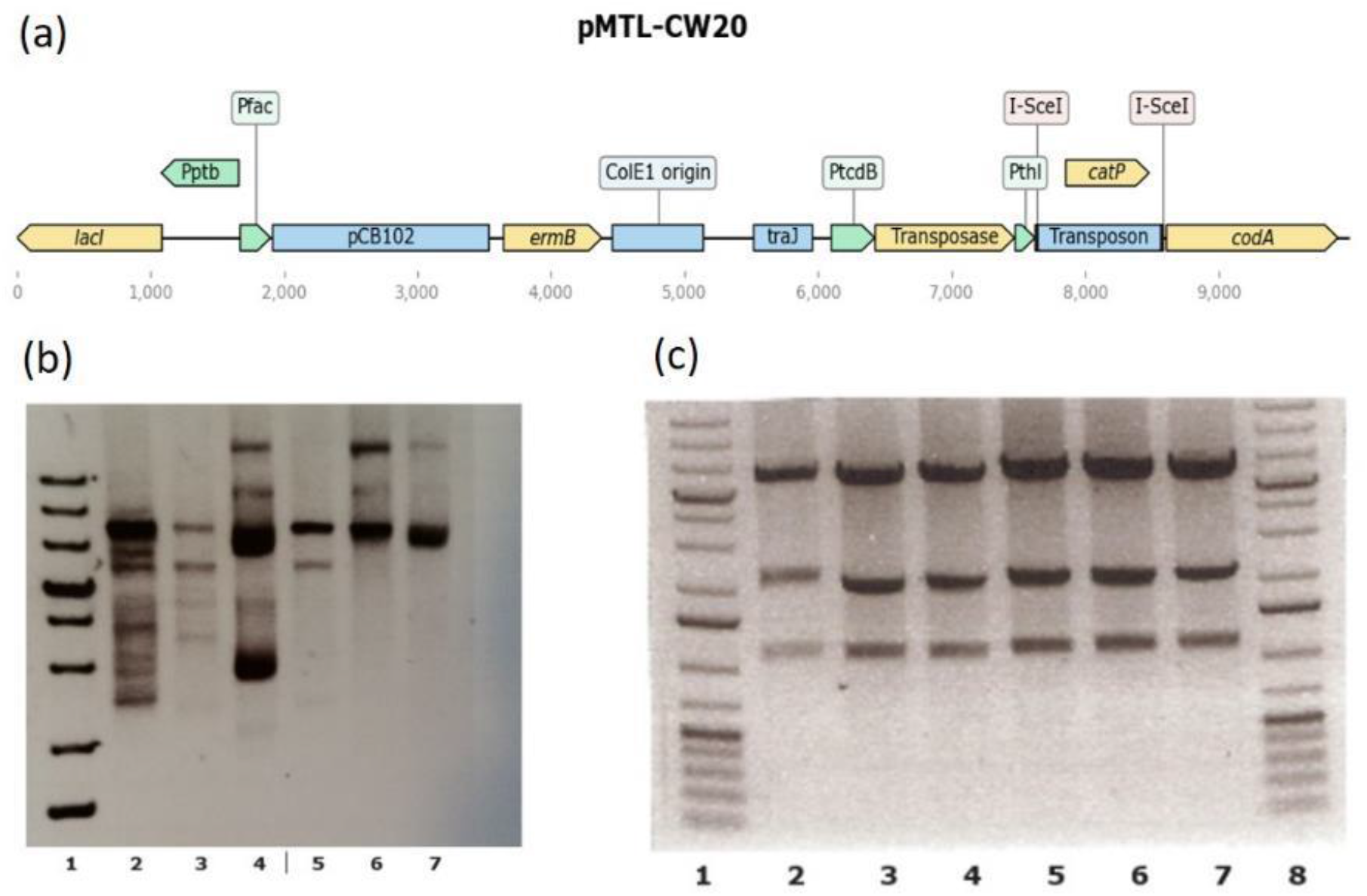
Transposon delivery plasmid pMTL-CW20. A) pMTL-CW20 is based on the pMTL-YZ14 plasmid described in ^24^ using components from the plasmid modular transfer series outlined in ^27^ as well as the *codA* from *E. coli* and I-SceI recognition sites. Replication occurs in *E. coli* via the pUC ColE1 origin of replication and the plasmid can be transferred to clostridial recipients using the *oriT* from RK2 ^28^. In clostridial hosts the plasmid is conditionally replicative where the presence of IPTG is the non-permissive condition. Transposition is achieved via a hyperactive Himar1 variant ^29^ which mobilises a mini-transposon containing the *catP* gene which confers chloramphenicol and thiamphenicol resistance. B) A verified pMTL-YZ14 plasmid was used to transform *E. coli* Top10 and transformant colonies used to inoculate overnight cultures. Plasmids prepared from overnight cultures were extracted and treated with SbfI. Movement of the transposon into various other parts of the vector was found to have occurred (2, 4, 6, and 7) while only lanes 3 and 5 exhibited the expected band pattern. ThermoFisher 1kb+ plus ladder is in lane 1 C) An analogous procedure using EcoRV was later followed using pMTL-CW20 instead of pMTL-YZ14. In this case all six plasmids exhibited the expected band pattern (lanes 2-7) with ThermoFisher 1kb+ ladder in lanes 1 and 8.

A second feature of pMTL-CW20 designed to control unwanted transposition was based on the provision of a promoter-less copy of the *E. coli codA* gene encoding cytosine deaminase to prevent premature transposition in the donor strain. Use of the previously described transposon-delivery vector pMTL-YZ14 ^24^ was characterised by inconsistent frequencies of transfer to the clostridial recipient and/or to variation in the effectiveness with which transposon mutants were generated once in *C. autoethanogenum*. These inconsistencies appeared to correlate with spontaneous plasmid rearrangements in the donor, as evidenced by unexpected DNA fragment profiles on agarose gels of diagnostic digests of the isolated plasmid DNA (Fig. 3). This was assumed to be due to transposition of the mini transposon from pMTL-YZ14 while in *E. coli* either into the genome or, as transposition into closed circular autonomous plasmids is preferred, into alternative positions in the vector backbone. The cut and paste nature of the transposition event would mean that plasmids would be generated that either no longer carried a mini-transposon or which had been affected in their maintenance or ability to transfer. Similar instabilities have been noted elsewhere ^25^.

Cytosine deaminase catalyses conversion of 5-fluorocytosine (5-FC) to the toxic product 5-fluorouracil (5-FU) which ultimately blocks DNA and protein synthesis. On the plasmid pMTL-CW20, *codA* is separated from its P_*thl*_ promoter (derived from the thiolase gene of *Clostridium acetobutylicum*) by the *catP* mini-transposon. Excision of the mini-transposon as a consequence of its transposition leads to expression of *codA*, a lethal event in the presence of exogenously supplied 5-FC. The addition of this feature to pMTL-CW20 improved the reproducibility with which the plasmid was transferred to *C. autoethanogenum* and appeared to prevent the occurrence of plasmid rearrangements (Fig. 3).

### Removal of plasmid-based reads

Suicide vectors are inappropriate for *C. autoethanogenum* since the low DNA-transfer rate means an unfeasible number of conjugations would need to be performed with a preliminary experiment resulting in five mutants from three conjugations. To compensate for this, a conditional replicon was utilised which has been described previously ^24^. To further remove any residual plasmid from the sequencing library I-SceI recognition sites were incorporated into pMTL-CW20 which provided a mechanism for removal of plasmid reads at the sequencing library stage. After adapter-ligation an I-SceI restriction is used to cleave the site between the adapter and the library primer binding site on the transposon, making those fragments originating from plasmid DNA unsuitable templates for the subsequent PCR amplification step as described in a similar strategy ^26^. In the initial transposon library grown on rich medium, 0.2% of reads mapped to the transposon-delivery plasmid. This compares favourably with a study on *Clostridiodes difficile* which used a replicative vector where 48% of the reads in the initial rich medium library mapped to the delivery plasmid ^6^.

### Generation of transposon library and growth in autotrophic conditions

Approximately 1.3 million colonies were pooled from 125 transposon selection plates and inoculated into 200 mL of rich medium (YTF) supplemented with thiamphenicol and IPTG. After 24 h of growth, genomic DNA was extracted from this culture and -80°C freezer stocks were made. This first genomic DNA extraction was used to determine the required gene set for growth on rich media, 100,065 unique insertion sites were found from this sample. Subsequently the freezer stocks were used to restore the mutant pool into a defined medium (PETC) supplemented with pyruvate as the carbon source. The PETC culture was used to inoculate a 1.5 L bioreactor containing fermentation medium which lacked a carbon source. The sole carbon and energy source after the inoculation was provided by CO gas sparged into the bioreactor with a gradual increase of CO. The pyruvate was quickly used up as shown by HPLC data (Supplementary Table S3) and *C. autoethanogenum* instead relied upon fixation of CO. The PETC medium provided no supplementary amino acids and instead relied on the native biosynthesis pathways of *C. autoethanogenum*. Vitamin requirements were met via the addition of Wolfes’ vitamin solution.

Samples for HPLC analysis of metabolites and for possible genomic DNA extraction and TIS analysis were taken on a daily basis. Samples from 72, 144, 168, 336, and 360 h of growth were used for sequencing. These sequencing data were used to determine the required gene set for growth using CO in a defined medium. Ultimately, the samples from 336 h and 360 h were used to determine the gene set required for growth on CO, these represent the endpoint of the reactor fed batch culture. The reactor endpoint was sequenced revealing 66,524 unique insertion sites.

### Functions of essential genes in heterotrophic compared with autotrophic conditions

The functions of candidate essential genes for growth in rich medium and minimal medium with CO as the carbon and energy source were compared using the KEGG database as summarised in Table 1. There were 439 genes (11%) identified as candidate essential genes out of a total of 4059 genes in the genome for heterotrophic growth on the rich medium YTF where fructose and yeast extract serves as a carbon and energy source (Supplementary Table S1). This is comparable with the number of genes in the Syn3.0 genome and close to the 404 reported in *Clostridiodes difficile* ^1,6^. As expected, genes involved in fundamental biological processes such transcription, translation, DNA replication and cell division are common in the rich media essential gene list. Eighteen of the twenty common amino acids have clearly annotated tRNA synthetases which appear essential except for tyrosine and asparagine. Tyrosine appears to exhibit redundancy via CLAU_1290 (*tyrZ)* and CLAU_1635. There is only one annotated asparagine tRNA synthetase *(asnB)* but it seems likely that there is another present (CLAU_2687) and that together they provide functional redundancy meaning that both genes are found to be non-essential. CLAU_2687 is currently annotated as a tRNA synthetase class II but is most likely to be an asparagine-specific tRNA synthetase. Another explanation for the non-essential status of the asparagine tRNA synthetase could be that *C. autoethanogenum* uses a mechanism common to many bacterial and archaeal taxa which entirely lack an asparagine tRNA synthetase. These taxa rely on a non-discriminating aspartic acid tRNA synthetase followed by an amidotransferase to generate asparagine-tRNAs ^30^.

**Table 1.** Functions of essential genes. Number of *C. autoethanogenum* essential genes for various KEGG functional categories on the rich medium YTF and the minimum fermentation medium with CO as a carbon and energy source

| Function | Rich | Minimal Medium + CO |
| --- | --- | --- |
| Metabolic Pathways | 98 | 156 |
| Ribosome | 38 | 42 |
| Microbial metabolism in diverse environments | 33 | 41 |
| Carbon metabolism | 25 | 30 |
| Aminoacyl-tRNA biosynthesis | 21 | 23 |
| Biosynthesis of amino acids | 18 | 36 |
| Hypothetical | 13 | 24 |
| Peptidoglycan biosynthesis | 11 | 12 |
| Pyruvate metabolism | 9 | 11 |
| Carbon fixation pathways in prokaryotes | 9 | 13 |
| DNA replication | 9 | 11 |
| Amino sugar and nucleotide sugar metabolism | 9 | 9 |
| Oxidative phosphorylation | 7 | 10 |
| Fatty acid metabolism | 6 | 7 |
| Fatty acid biosynthesis | 6 | 7 |
| Propanoate metabolism | 6 | 7 |
| Quorum sensing | 5 | 7 |
| Protein export | 5 | 9 |
| Purine metabolism | 5 | 14 |
| Bacterial secretion system | 5 | 9 |
| RNA degradation | 5 | 7 |
| RNA polymerase | 4 | 4 |
| Pyrimidine metabolism | 4 | 10 |
| Citrate cycle (TCA cycle) | 3 | 7 |
| Phosphotransferase system (PTS) | 2 | 2 |
| Folate biosynthesis | 2 | 5 |
| Biotin metabolism | 1 | 2 |
| Flagellar assembly | 1 | 1 |
| Bacterial chemotaxis | 1 | 2 |

The candidate essential gene list for rich medium calls into question several of the annotations in the *C. autoethanogenum* genome. For instance, CLAU_0265 which is annotated as a small acid-soluble spore protein is required on rich medium despite *C. autoethanogenum* C24 never having been observed to sporulate. In addition, sporulation should never have been required in the library preparation process. The gene must, therefore, have an additional or alternative role. Much functional genomics work has yet to be performed on *C. autoethanogenum* since there are 44 rich medium essential genes annotated as hypothetical proteins (Supplementary Table S1).

In total, 758 genes (19% of the genome) were predicted to be required for autotrophic growth by the endpoint of the CO-fed reactor (Supplementary Table S1). This includes all of the ‘core’ gene set which were also required on rich medium and all of the genes required to grow on minimal medium lacking amino acids. The core gene set was predicted to be comprised of 439 genes. This means that 319 genes are likely to be required for the synthesis of all amino acids and utilisation of CO as a carbon and energy source. As vitamins were provided, their biosynthetic pathways were not expected to be represented, similarly nitrogen and sulphur were supplied in the medium. All genes and their predicted essentiality status in each experimental condition are presented in Supplementary Table S1. Comparing two predicted required gene lists from different times in an experiment is an imperfect method of deducing condition-specific genes, but the data is nevertheless extremely informative.

### Essential genes of the Wood-Ljungdahl (WLP) pathway

In order to grow using CO as a sole carbon and energy source it is necessary for *C. autoethanogenum* to use two molecules of CO to form one molecule of acetyl-CoA. Acetyl-CoA consists of a methyl group, a carbonyl group and the CoA cofactor. The methyl group is supplied by the action of the bifunctional CODH enzyme which oxidises CO to CO_2_, this CO_2_ molecule then follows the Eastern branch of the WLP before being combined with another CO molecule and CoA by the acetyl-CoA synthase (ACS). It was therefore expected that all the genes involved in the WLP would be required for growth on CO. A complete WLP was indeed found in the list of essential genes and has been mapped out in Fig. 4. The WLP was not required during heterotrophic growth despite the fact that it is utilised during heterotrophic growth to fix CO_2_ released during glycolysis using the reducing equivalents generated by glycolysis ^23^.

**Figure 4.**
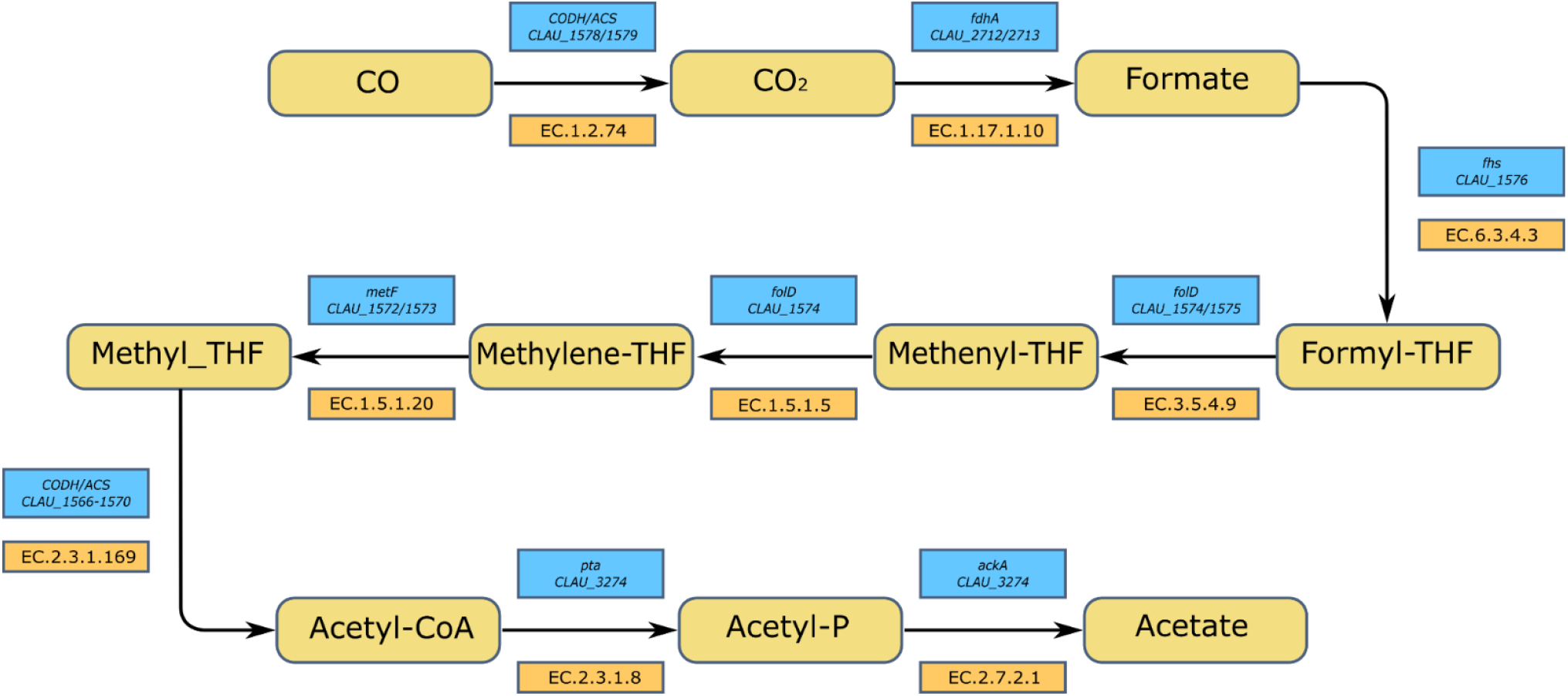
Essential Genes of the Wood-Ljungdahl pathway. Route from CO to acetate showing the expected gene/locus tag for each step. Each of the locus tags listed was required for growth on CO.

To generate ATP, *C. autoethanogenum* is reliant on generating a transmembrane electrochemical gradient via the intrinsically important ^31^ Rnf complex and a membrane integral ATP synthase ^18, 32^. The Rnf complex of *C. autoethanogenum* is encoded by the region CLAU_3144-CLAU_3149. A comparison of the insertion sites found in heterotrophic and autotrophic conditions for this region is shown in Fig. 5. With exception of *rnfB*, all of the encompassed genes were found to be essential for growth on CO (Fig.5), confirming previous observations that inactivation of these genes in either *C. ljungdahlii* and *Acetobacterium woodii* curtailed growth on H_2_ + CO_2_ ^33,34^. Despite *rnfB* being above the insertion index threshold for essentiality on CO, it is significantly under-represented when the data obtained from cells grown on pyruvate is compared with CO (log_2_ fold change = -2.86, p-value = 1.16E-11). It may be the case that *rnfB* encodes a non-essential component of the complex which aids functionality but is not required for it. In Methanogens RnfB has been characterized as an entry point for electrons to the Rnf complex ^35^.

**Figure 5:**
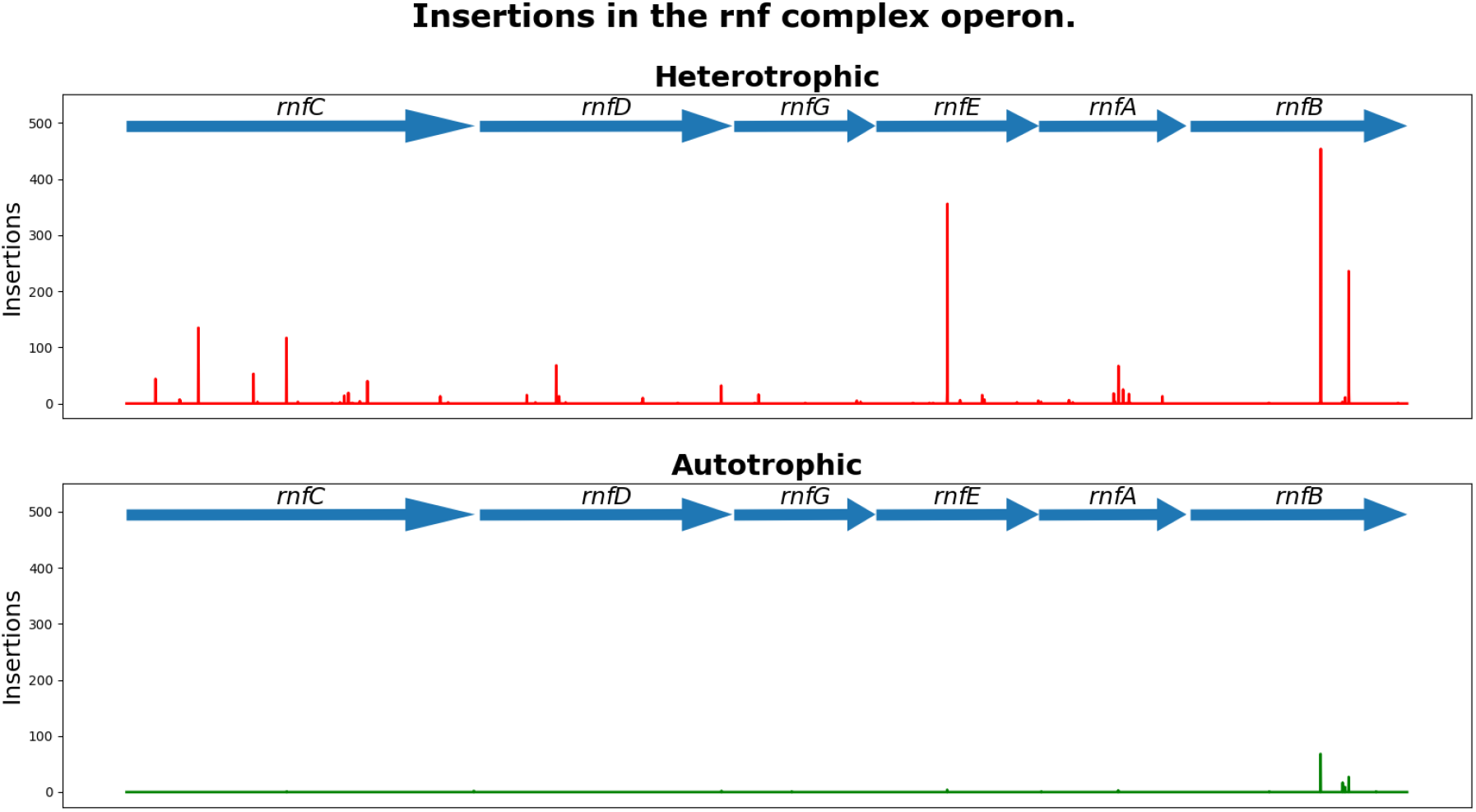
Insertions in the Rnf complex region. Number of reads detected along the genomic region encoding the Rnf complex for heterotrophic and autotrophic conditions. Insertions are relatively abundant in heterotrophic conditions implying importance for the complex under autotrophic conditions.

### The importance of Nfn for autotrophic growth

In order to further verify the calling of gene essentially under specific conditions using our parameters, a candidate gene was selected for directed CRISPR mutagenesis. The *nfn* gene (CLAU_1539) encodes an electron-bifurcating ferredoxin-dependent transhydrogenase, responsible for the production of NADPH from NADH and Fd^2-^, thus recycling NAD+. Our TIS data analysis found that the *nfn* gene was non-essential when *C. autoethanogenum* was grown on rich medium or when grown on minimal medium with pyruvate, but when autotrophic conditions were used the gene was essential. This suggested that a directed CRISPR knockout mutant should be obtainable while the culture is grown under heterotrophic conditions but should fail to survive when transferred to autotrophic conditions. A CRISPR in-frame deletion mutant of *nfn* (Δ*nfn*) was created which was viable on rich media, and on minimal medium with pyruvate as a carbon source, but was unable to grow when CO was the sole carbon and energy source.

Initially the Δ*nfn* strain was characterised in serum bottles, using minimal PETC media and either 10 mM of sodium pyruvate, or 1 bar of CO in the headspace, as the carbon and energy source. Serum bottles were inoculated with 1 ml (1:50 inoculum) of a late exponential culture grown in the anaerobic cabinet on minimal media with fructose as a carbon source. The cultures grown on pyruvate grew similarly to the wild type control, however, no evidence of growth was evident when CO was used instead of pyruvate as the carbon and energy source. This inability to utilise CO as a carbon and energy source was further demonstrated on a larger scale using a fed-batch CSTR experiment, whereby a 1.5 L culture was inoculated with 150 ml of an early exponential culture grown on minimal media and pyruvate. The pH was controlled with NaOH and H_2_SO_4_, and sparged through continual addition of nitrogen at a rate of 60 ml/min. At the time of inoculation 5 mM of sodium pyruvate was added to the culture. Once an OD_600_ of approximately 0.3 had been reached, CO was introduced at a rate of 10 ml/min. In the case of the wild type culture, the strain was able to adapt to the CO carbon and energy source and after 48 h the OD continued to increase after the pyruvate had been depleted. In the case of the *nfn* mutant, the culture was not able to adapt to utilising CO, and the optical density rapidly declined following depletion of the pyruvate.

### Assessment of metabolic modelling-derived gene essentiality

Experimentally confirmed gene essentiality for growth on minimal medium supplemented with CO was compared against the predicted essentiality calculated from a metabolic model of *C. autoethanogenum* by means of Flux Balance Analysis (FBA) ^36,37^ (Supplementary Table S2). To that end, the confusion matrix was generated (Table 2) to calculate Matthew’s correlation coefficient (MCC), a robust metric ranging from -1 to +1 used to evaluate binary classifications (such as essential or non-essential gene) ^38^. A MCC = 0.40 was obtained, indicating that the model has a certain predictive capability. Interestingly, a value of 0.69 has been reported for a similar study in *E. coli* ^39^. Considering the high level of curation of models for this bacterium compared to recently published ones ^36,40^, it is safe to assume that result to be an upper achievable value.

**Table 2.** Confusion matrix for gene essentiality comparison. TP = true positive, FP = false positive, FN = false negative, and TN = true negative. A perfect correlation (MCC = +1) would require TP = 179, FP = 0, FN = 0, and TN = 353.

| Predicted | Experimental |  |
| --- | --- | --- |
|  | Essential | Non-essential |
| Essential | TP = 110 | FP = 76 |
| Non-essential | FN = 69 | TN = 277 |

## DISCUSSION

The successful application of TIS to *C. autoethanogenum* has provided a wealth of information on gene essentiality in this industrially important acetogen and represents the most thorough analysis of its kind performed to date in clostridia. The essentiality status of all *C. autoethanogenum* genes can now be consulted (in Supplementary Table S1) before directed knockouts are made. The conditionally essential genes are particularly useful when hypothesising on gene functions and may be the starting point for further research with the 30 hypothetical proteins which were not required on rich medium, but were essential for minimal medium autotrophic growth being the most obvious examples.

### Essentiality in the Wood-Ljungdahl pathway (WLP)

The generation of the methyl group from CO first requires its oxidation to CO_2_ by a CODH. *C. autoethanogenum* possesses three such enzymes genome, namely CLAU_1578/CLAU_1579 (*acsA*), CLAU_2924 (CAETHG_3005) (*cooS1*) and CLAU_3807 (CAETHG_3899) (*cooS2*) ^17^. Their independent interruption via ClosTron mutagenesis ^41^ showed that autotrophic growth was only abolished in the *acsA* knockout mutant suggesting it is the only CODH required for growth on either CO or CO_2_. Our data validates this conclusion with only *acsA* being predicted as required for growth on CO but not on YTF. The remaining putative CODH genes were required in neither condition and there was no substantial change in insertion index between YTF and CO conditions. The conditionally essential CODH encoded by *acsA* has an internally translated stop codon (TGA) not found in the equivalent genes of related organisms and can alternatively be thought of as two ORFs (CLAU_1578 and CLAU_1579) although it appears that CLAU_1579 makes no separate product. It has been demonstrated that *acsA* can be translated either as a 44 kDa protein or as a 69 kDa protein depending on whether the TGA internal stop codon is the end of translation or whether it causes the incorporation of a selenocysteine residue ^23^. It appears from our data that both ORFs are required under autotrophic conditions. Thus, the 44 kDa protein alone does not appear to be sufficient for autotrophy and cells apparently require the 69 kDa protein to be autotrophic.

There are three putative formate dehydrogenases in the *C. autoethanogenum* genome encoded by CLAU_0081, CLAU_2712/CLAU_2713 (*fdhA*), and CLAU_2907. Of these, *fdhA* alone appears to be essential only on CO while the remaining two genes are required in neither tested condition. The most important formate dehydrogenase is therefore *fdhA* which is found in a complex with an NADP-specific electron-bifurcating [FeFe]-hydrogenase (Hyt) ^42^. Two of the three putative formate dehydrogenases are selenoenzymes which may be higher efficiency than the cysteine-containing analogues, it is therefore tempting to speculate that the non-selenoenzyme formate dehydrogenase may be present as a backup for low selenium conditions ^43^. However, it appears from our data that neither CLAU_0081 nor CLAU_2907 could provide sufficient activity in the *fdhA* mutants for them to not be outcompeted causing *fdhA* to appear essential under autotrophic conditions.

The steps from formate to methyl-THF are catalysed by the products of CLAU_1572-CLAU_1576 which all appear to be required for growth on CO. CLAU_1574 and CLAU_1576 additionally appear to also be required for growth on the rich medium.

The methyl group of methyl-THF is transferred to the Corrinoid Iron-Sulfur Protein (CoFeSp) cofactor before being combined with the carbonyl group supplied by another molecule of CO by the action of the ACS (acetyl-CoA synthase). The ACS is encoded by the region CLAU_1566-70 in which CLAU_1566, CLAU_1568, and CLAU_1569 were only required on CO whereas CLAU_1567 and CLAU_1570 were required on CO and on YTF.

### Essentiality in the metabolism from acetyl-CoA

There are four main carbon compounds at the end points of metabolism for *C. autoethanogenum*: acetate, ethanol, 2,3 – butanediol, and lactate. The route to acetate from acetyl-CoA proceeds through acetyl phosphate; catalysed by the enzymes phosphate acetyltransferase (encoded by *pta*) and acetate kinase (encoded by *ackA*). Both *pta* and *ackA* were found to be essential when growing on rich medium. The step from acetyl phosphate to acetate regenerates ATP and so this pathway may be required for energy generation.

However, *pta* has been knocked out in *C. ljungdahlii* ^44-46^ where it significantly impaired growth rates and acetate formation. The *C. ljungdahlii pta* knockout may be viable only because of a second putative phosphate acetyltransferase (WP_063556670.1) which is annotated as a bifunctional enoyl-CoA hydratase/phosphate acetyltransferase and has no homolog in *C. autoethanogenum*. The bifunctional enoyl-CoA hydratase/phosphate acetyltransferase may be producing sufficient ATP for cells to be viable. The absence of an alternative phosphate acetyltransferase in *C. autoethanogenum* is likely the cause of the essentiality of *pta* in our data.

The route to ethanol can proceed from acetyl-CoA either straight to acetaldehyde and then to ethanol or via acetate, then acetaldehyde and finally ethanol. The more direct route from acetyl-CoA to acetaldehyde is catalysed by an acetaldehyde dehydrogenase (EC 1.2.1.10) which could be encoded by an estimated five genes within the *C. autoethanogenum* genome (CLAU_1772, CLAU_1783, CLAU_3204, CLAU_3655, CLAU_3656) none of which appear to be required in either growth condition. This could represent redundancy between these genes which further knockout studies could aim to confirm, or it could be that this route to ethanol is not required. The alternative route to ethanol via acetate is similar in that there are two predicted genes (CLAU_0089 and CLAU_0099) which could encode an aldehyde ferredoxin oxidoreductase (AOR; EC 1.2.7.5) but neither of them appear to be essential in either growth condition. In case this the result is best explained by a lack of biological necessity for this reaction since it has been shown that a double AOR knockout strain was still viable autrophically ^23^.

There are two candidate genes encoding pyruvate synthase enzymes for formation of pyruvate from acetyl-CoA (CLAU_0896 and CLAU_2947) of which only CLAU_2947 appears to be required; this is true in both growth conditions. All of the genes encoding functions for the pathways leading to lactate and 2,3-butanediol appear non-essential. In the case of the conversion of acetolactate to acetoin and in the production of lactate utilising NADH there appears to be only one gene encoding the relevant functions (CLAU_2851 and CLAU_1108 respectively); in these cases redundancy is unlikely to be the reason for their non-essential status meaning it is more likely these are unnecessary biological routes.

Overall, our findings highlight that TIS represents a powerful functional genomics tool which can be applied to less genetically tractable organisms using the methods applied here. Presented data allows a confident determination of the Wood-Ljungdahl pathway genes of *C. autoethanogenum* and opens up future avenues of investigation into the genes which are essential for autotrophic growth with no obvious reason as to why.

## MATERIALS AND METHODS

### Microbiology

*E. coli* DH5alpha (NEB) was used for all for cloning and sExpress ^47^ as a conjugal donor. Strains were cultured at 37 °C in LB broth with appropriate antibiotic supplementation and 5-FC in a shaking incubator or on LB agar in a static incubator. *C. autoethanogenum* was cultured and manipulated in an anaerobic workstation (Don Whitley) with an atmosphere of 80% nitrogen, 10% carbon dioxide and 10% hydrogen at 37 °C. The three media used, were YTF (Table S4-S7), PETC (Table S8-S10) and Fermentation (Table S11-S13) medium. Plasmids were transferred from sExpress to *C. autoethanogenum* as detailed in Woods et al., 2019 ^47^. Briefly, this involved mixture of the donor and recipient cultures together and incubation on a non-selective YTF plate for 20 h before harvesting and plating onto selective YTF agar. Antibiotic selection for transposon plasmids was performed using chloramphenicol (25 µg/ml) and erythromycin (500 µg/ml) in *E. coli* or thiamphenicol (15 µg/ml) and clarithromycin (6 µg/ml) in *C. autoethanogenum*. Kanamycin (50 µg/ml) was used to select for the sExpress donor strain. D-cycloserine (250 µg/ml) was used to counter-select the sExpress donor strain. Fluorocytosine (FC) was supplemented at 30 µg/ml and IPTG at a concentration of 1 mM. Plasmid pMTL-CW20 maybe sourced from www.plasmidvectors.com.

### DNA manipulations

Genomic DNA purifications were performed using bacterial gDNA extraction kits from Sigma Aldrich. Plasmid DNA was purified with mini-prep kits from NEB. Screening PCRs were performed using DreamTaq polymerase (Thermo Fisher). Oligonucleotides were synthesised by Sigma Aldrich. Sanger sequencing was performed by Source Bioscience.

### Mutant generation using CRISPR

A CRISPR in-frame deletion vector was designed as previously described using the pMTL40000 CRISPR vector series ^48^. In this case we employ the trCas9 nickase variant under control of the *fdx* promoter from *C. sporogenes*, a unique sgRNA under control of the constitutive *araE* promoter from *C. acetobutylicum* targeting *nfn*, and a homologous recombination cassette to allow the precise in-frame deletion of *nfn*. Following vector assembly, the construct was transferred to wild type *C. autoethanogenum* by conjugation using sExpress as the *E. coli* donor strain ^47^. Following two rounds of selection on thiamphenicol and D-cycloserine, to select for recipient strains harbouring the CRISPR vector and counterselect the *E. coli* donor cells, respectively, a colony PCR screen was performed on resultant colonies, amplifying from the genomic locus flanking the regions selected for homologous recombination. The screen revealed that the *nfn* knockout mutant was indeed present in the population, and the strain was sub-cultured for storage and preparation of genomic DNA. Sanger sequencing from a high-fidelity PCR product confirmed the precise in-frame deletion of *nfn*.

### Assessment of transposon vectors

Transposon delivery vectors were transferred to *C. autoethanogenum* via conjugal transfer from sExpress and selected for on YTF agar plates supplemented with clarithromycin and D-cycloserine. Colonies were harvested from selection plates by flooding with PBS and the entire cell suspension was serially diluted and spread onto YTF agar plates supplemented with either clarithromycin and IPTG, thiamphenicol and IPTG or clarithromycin to determine the transposition frequency and plasmid-retention in the presence of IPTG.

### Transposon library creation

The transposon-delivery vector pMTL-CW20 was transformed into an *E. coli* conjugative donor strain sExpress ^47^ which was used to transfer the plasmid into *C. autoethanogenum* C24. Twelve conjugations were performed simultaneously producing a total of around 81,000 transconjugant colonies on YTF agar supplemented with D-cycloserine and clarithromycin. All transconjugants were pooled and plated onto agar plates supplemented with IPTG, lactose and thiamphenicol to select for transposon mutants. Transposon mutant colonies were then harvested and inoculated to YTF broth supplemented with IPTG and thiamphenicol. The rich medium sequencing samples were taken from this liquid phase which was used to inoculate PETC pyruvate medium. The PETC pyruvate culture was allowed to reach stationary phase before being used as inoculum for the bioreactor where CO served as the carbon and energy source; a DNA sample was taken from PETC pyruvate at the point of bioreactor inoculation. Samples were taken from the bioreactor to check for the presence of pyruvate, monitor the OD and to serve as DNA samples for identification of insertion sites.

### Bioinformatics and metabolic modelling

Experimentally confirmed essential genes were compared against metabolic modelling-derived essentiality. Lists of essential genes were generated using the BioTraDIS toolkit approach as previous described ^49^. A genome-scale model (GSM) of CO-fed *C. autoethanogenum* was handled using the COBRA Toolbox in MATLAB R2017b to predict gene essentiality ^36,50^. Briefly, the wild type model was subjected to Flux Balance Analysis (FBA) by selecting the maximization of the biomass yield as the objective function ^37^. Simulated single gene deletions provoking a reduction of at least 5% of the optimal specific growth rate of the wild type were deemed essential. Finally, Matthew’s correlation coefficient was used as a metric to assess the quality of the GSM predictions, where “1” is a perfect correlation between experimental and predicted gene essentiality, “0” no correlation, and “-1” perfect anti-correlation ^39^. The model and the scripts are available in GitHub (https://github.com/SBRCNottingham/C.auto_essentiality).

Sequencing and bioinformatics. Sequencing library preparation was performed as an amplicon library using a splinkerette adapter ^49^. Genomic DNA was fragmented to an average of 400 bp using a covaris sonicator followed by bead purification using NEB sample preparation beads at a ratio of 1.5X beads to sample. Fragmented DNA was end repaired and A-tailed using the NEB Ultra II library preparation kit. Splinkerette adapters were ligated onto the end of A-tailed fragments with reagents from the Ultra II library preparation kit. A 1X bead purification was performed before an I-SceI digest step to cleave plasmid DNA between the library primer and P7 primer. Another 1X bead purification was performed before PCR amplification of the transposon junctions using KAPA HiFi polymerase. An initial denaturing step of 95°C for two min was followed by 20 rounds of 95°C for 20 sec, 61°C for 30 sec then 72°C for 30 sec before a final extension of two min at 72°C was performed.

PCR products with a size range of 250-500 bp were gel extracted from a low-melt agarose gel using the NEB monarch gel extraction kit. Gel extracted products were analysed on an Agilent bioanalyser using a DNA 1000 chip and quantified via Qubit and qPCR. Two separate runs were performed on an Illumina MiSeq.

Raw sequences were trimmed before filtering for reads which contain the expected transposon tag. The transposon tag was removed from reads which could then be mapped to the *C. autoethanogenum* genome to identify insertion sites. The BioTraDIS analysis pipeline was used for these steps and for subsequent analysis ^49^.

## ACKNOWLEDGEMENTS

This work was funded by the Biotechnology and Biological Sciences Research Council [grant numbers BB/L502030/1, BB/K00283X/1, BB/L013940/1]. NPM acknowledges funding from LanzaTech as part of BB/L502030/1. The funders had no role in study design, data collection and analysis, decision to publish, or preparation of the manuscript. Sequencing was performed by Deep Seq (University of Nottingham) with thanks to Nadine Holmes, Matthew Carlile, and Victoria Wright.

## Author Contributions

CW, CMH and NPM planned the study. CW and CMH undertook the experimental work. CTA performed the metabolic modelling. Data analysis was undertaken by CW. CW and CMH drafted the manuscript. All authors reviewed, edited, and approved the final version of the manuscript.

